# Phylogeny and biogeography of *Ceiba* Mill. (Malvaceae, Bombacoideae)

**DOI:** 10.1101/2020.07.10.196238

**Authors:** Flávia Fonseca Pezzini, Kyle G. Dexter, Jefferson G. de Carvalho-Sobrinho, Catherine A. Kidner, James A. Nicholls, Luciano P. de Queiroz, R. Toby Pennington

## Abstract

The Neotropics is the most species-rich area in the world and the mechanisms that generated and maintain its biodiversity are still debated. This paper contributes to the debate by investigating the evolutionary and biogeographic history of the genus *Ceiba* Mill. (Malvaceae: Bombacoideae). *Ceiba* comprises 18 mostly neotropical species endemic to two major biomes, seasonally dry tropical forests (SDTFs) and rain forests and its species are one of the most characteristic elements of neotropical SDTF, one of the most threatened biomes in the tropics. Phylogenetic analyses of DNA sequence data from the nuclear ribosomal internal transcribed spacers (ITS) for 30 accessions representing 14 species of *Ceiba* recovered the genus as monophyletic and showed geographical and ecological structure in three main clades: (i) a rain forest lineage of nine accessions of *C. pentandra* sister to the remaining species; (ii) a highly supported clade composed of *C. schottii* and *C. aesculifolia* from Central American and Mexican SDTF plus two accessions of *C. samauma* from inter Andean valleys from Peru; and (iii) a highly supported South American SDTF clade including 10 species showing little sequence variation. Within this South American clade, no species represented by multiple accessions were resolved as monophyletic. We demonstrate that the patterns of species age, monophyly and geographical structure previously reported for SDTF species within the Leguminosae family are not shared by *Ceiba*, suggesting that further phylogenetic studies of unrelated groups are required.

**HIGHLIGHTS:**

- This paper provides a well sampled phylogeny of the iconic genus *Ceiba*, one of the most characteristic tree genera of neotropical Seasonally Dry Tropical Forest (SDTF).
- There is a clear phylogenetic signal for biome preference and geographic structure in *Ceiba*.
- We estimate a mid Miocene origin for *Ceiba*, with the stem node age of the genus estimated at 21.1 Ma and the crown node age at 12.7 Ma.
- *Ceiba* species have young stem ages in the SDTF clade but old stem ages in rain forest species.
- Patterns of species age, monophyly, ecological and geographical structure reported for SDTF species are only partially shared by *Ceiba*, an iconic genus of neotropical SDTF.

## INTRODUCTION

The Neotropics is the most species-rich region in the world and the mechanisms that generated and maintain its biodiversity are under constant discussion. Through evolutionary time, the Neotropics have been climatically and geologically dynamic, resulting in a diversity of biomes, from deserts to tropical rain forests (Hughes et al. 2013, Rangel et al. 2018). To understand the history and dynamics of those biomes, molecular phylogenetic and phylogeographic approaches have been used, because the relationships of taxa allow inferences to be made of the historical relationships amongst biomes and areas (Pennington et al. 2006). In recent years the dichotomy regarding the ‘cradle’ vs. ‘museum’ debate (Stebbins 1974) explaining neotropical diversity has given way to a more nuanced approach, considering that plant diversification patterns may be recent, old, slow or rapid, even within individual clades (Hughes et al. 2013, Koenen et al. 2015). As suggested in the literature more than 10 years ago (Wiens and Donoghue 2004, Pennington et al. 2006), this heterogeneity in diversification timing and rate within and among clades may be related not only to climatic and geological events, but also to the age and ecological differences of the biomes. For example, geologically old biomes (e.g., rain forest) are likely to have provided lineages that colonised newer biomes (e.g., savannas) and the relative difficulty of evolving adaptations such as drought tolerance or the ability to survive fire might determine whether a lineage can adapt to a new biome (niche evolution; Simon et al. 2009, Pennington & Lavin 2016) or remains confined to the same biome (niche conservatism) over evolutionary timescales (Crisp et al. 2009).

Clades endemic to two of the major Neotropical biomes, seasonally dry tropical forests (SDTFs) and rain forests, give good examples of different and distinctive phylogenetic and biogeographic patterns, suggesting an interaction of ecology and phylogeny over evolutionary timescales (Pennington et al. 2011, Pennington and Lavin 2016, Dexter et al. 2017). SDTFs occur on fertile soils and are characterized by the absence of fire adaptation in the flora and a predominantly continuous tree canopy, which becomes more open in the drier sites, with plants shedding up to 90–95% of their leaves during the five to six month long dry season (Murphy and Lugo 1986, Pennington et al. 2009). This biome has been one of the least studied, but is one of the most threatened in the tropics (Miles et al. 2006, DRYFLOR 2016). It occurs in disjunct areas throughout the Neotropics and has high beta-diversity and plant species endemism (Pennington et al., 2009; DRYFLOR, 2016). Leguminosae and Bignoniaceae are often the most species rich and dominant families in SDTF, but species from the Bombacoideae clade (Malvaceae), the subject of this paper, are often common and distinctive.

SDTF-confined clades contain species that often resolve as monophyletic and with old stem ages in DNA-sequence-based phylogenies (Pennington and Lavin 2016). In addition, the geographically structured phylogenetic pattern characteristic of clades in this biome suggests dispersal-limitation caused by the stable ecological conditions of the biome maintained over long evolutionary timescales (Pennington et al. 2010, Hughes et al. 2013). By contrast, tree clades confined to the Amazon rain forest, the largest expanse of tropical forest in the world, are suggested to contain more non-monophyletic species, more species with young stem ages and clades that lack geographical phylogenetic structure (Dexter et al. 2017). These rain forest patterns might be explained by frequent dispersal and subsequent successful colonization over evolutionary timescales (Pennington & Lavin, 2015) (Fig. 1).

**Figure 1.**
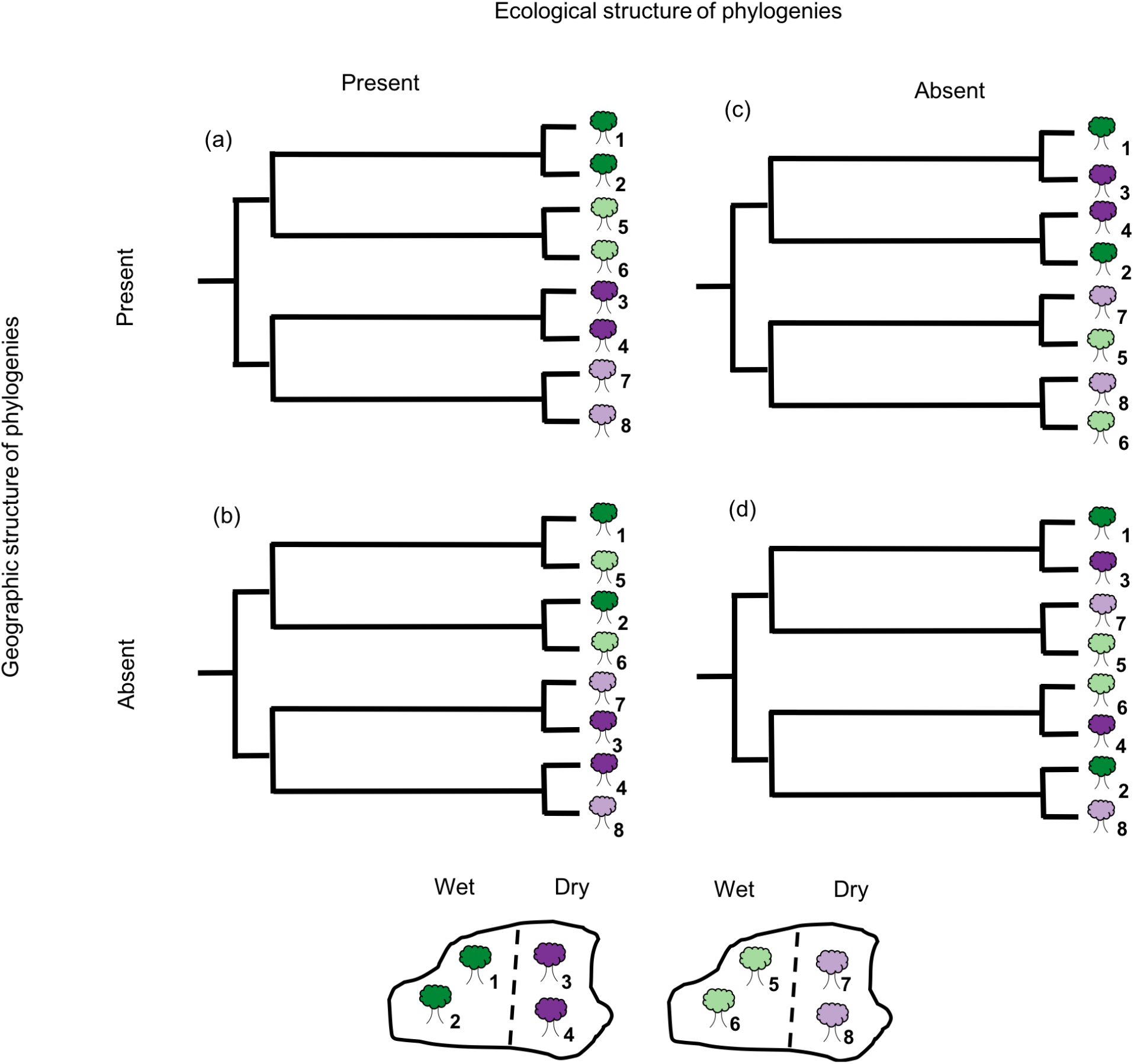
Two hypothetical islands, each with an area of seasonally dry tropical forest and rain forest. In total eight different species occur in the two biomes and the different islands and are represented by different colours (dark green species 1, dark green species 2, dark purple 3, etc.). Hypothetical phylogenies showing patterns of presence or absence of geographical and ecological structures (after Graham & Fine 2008).

The neotropical genus *Ceiba* Mill. (Malvaceae: Bombacoideae) comprises 18 species grouped into taxonomic sections *Ceiba* and *Campylanthera* (Schott & Endl.) K. Schum. based on morphological characters of pollen and staminal appendages. It is one of the most characteristic elements of many neotropical SDTFs. However, it also contains species confined to the Amazon rain forest and is therefore a good case study to investigate biome-specific differences in the nature of species and their diversification trajectories.

*Ceiba* species have digitate leaves, aculeate spines on the trunk and branches and can vary from 50 m canopy emergents in seasonally flooded várzea forests in the Amazon (*C. pentandra*) to 2 m treelets on rocky outcrops (campos rupestres) in Minas Gerais, Brazil (*C. jasminodora*). In some species (*C. chodatii*, *C. pubiflora*, *C. glaziovii, C. speciosa*) the trunk can be ventricose (swollen), explaining its vernacular names barriguda (“swollen belly”; Brazil) and palo borracho (“drunken tree”; Peru). Most species are deciduous and flower when leafless. They occur mostly in SDTF, with the exception of the widespread *C. samauma*, *C. speciosa* and *C. pentandra* that also occur in more humid environments, and *C. lupuna*, which is the only species restricted to rain forests (Fig. 2). On average, each of the thirteen SDTF species have a narrower geographical distribution when compared to the five rain-forest-inhabiting species (Fig. 2).

**Figure 2.**
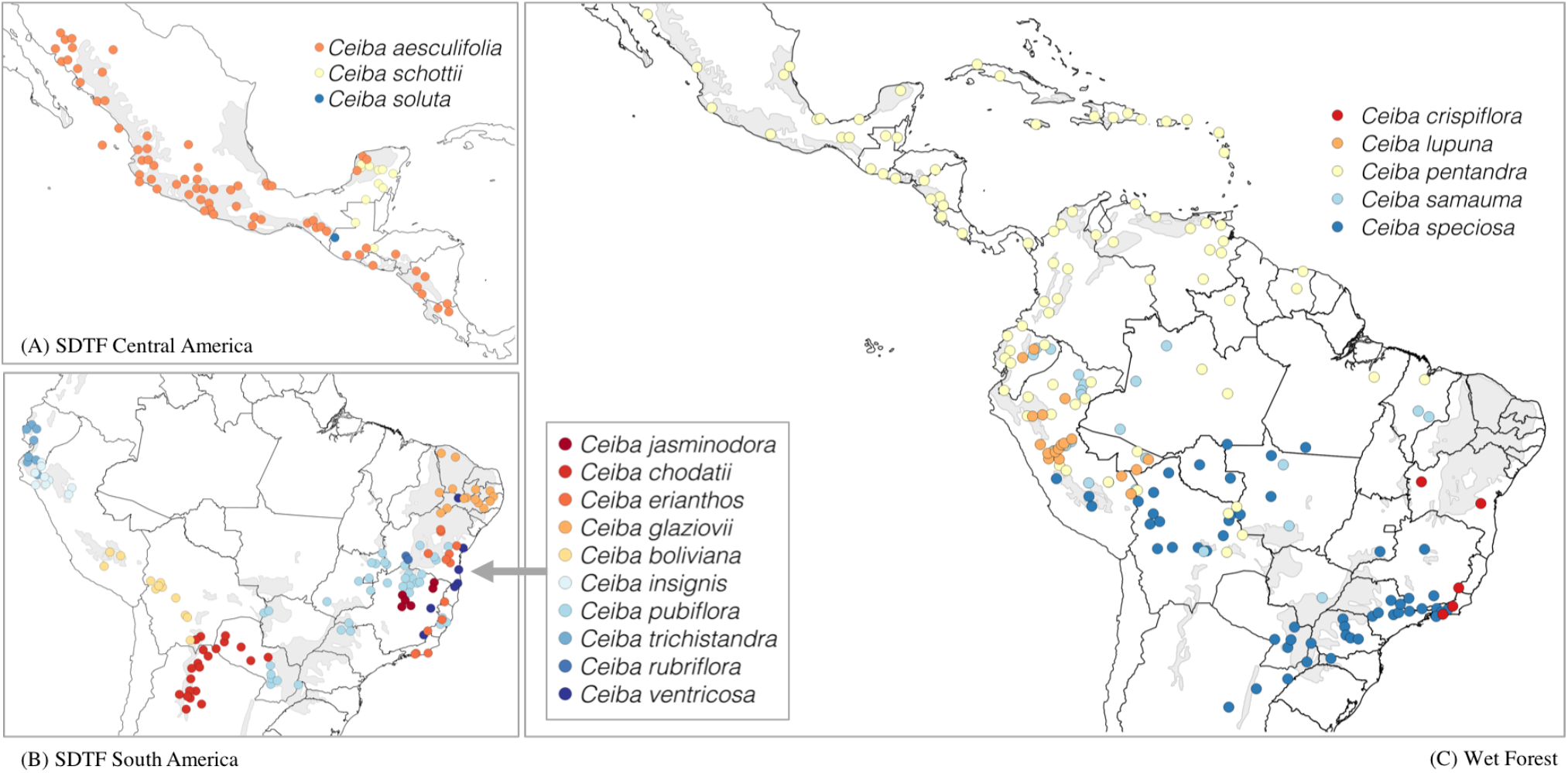
Distribution of 18 *Ceiba* species in three maps: (A) three species from SDTFs in Central America and North America, (B) ten species from SDTFs in South America and (C) five species from rain forests from Latin America. Grey areas represent the distribution of SDTF following DRYFLOR (2016). Occurrence records adapted from Gibbs and Semir (2003).

Previous Bayesian analyses of sequence data from the nuclear ribosomal internal and external transcribed spacers (ITS and ETS) and plastid markers (matK, trnL-F, trnS-trnG) for 13 species recovered *Ceiba* as monophyletic and sister to *Neobuchia paullinae* (Duarte et al. 2011, Carvalho-Sobrinho et al. 2016). Together with *Spirotheca*, *Pochota fendleri* sensu Alverson and Duarte (2015), and *Pseudobombax*, these taxa form the well supported “striated bark” clade (Carvalho-Sobrinho et al. 2016). However, relationships within *Ceiba* were poorly resolved and only one individual per species was included in the phylogeny. *Ceiba* has a historically complex taxonomy with species boundaries still confused, which is aggravated by the fact that herbarium specimens are often incomplete because individuals produce flowers and fruits when leafless. Therefore, a well sampled phylogeny with multiple accessions per species could be a useful tool to explore the nature of species in *Ceiba*.

This paper investigates the evolutionary history of *Ceiba* species. We aim to assess whether the *Ceiba* phylogeny is geographically or ecologically structured and if species confined to SDTFs are resolved differently in the phylogeny as compared with rain forest species (i.e., monophyletic on long stem lineages).

## METHODS

### Taxon sampling

We present the best sampled phylogeny of the genus *Ceiba* to date, covering 30 accessions representing 14 of the 18 species described for the genus. Critically, this study is the first to sample multiple individuals per species for six species (Table S1). As outgroups, we included 10 accessions representing species of the closest sister clades (Carvalho-Sobrinho et al. 2016): *Pseudobombax*, *Spirotheca, Eriotheca*, and *Pochota fendleri*. The full data set represents a combination of new sequence data from field surveys as well as from herbarium specimens, doubling the number of accessions of *Ceiba* in relation to the previous study by Carvalho-Sobrinho et al. (2016).

### DNA sequence data

We used the ITS region to investigate species relationships in *Ceiba*. In Bombacoideae, this region has been widely explored to help elucidate relationships among genera and species (Baum et al. 1998, Duarte et al. 2011, Carvalho-Sobrinho et al. 2016), and to investigate genetic structure among populations (Dick et al. 2007). Previous systematic studies in Bombacoideae used a combination of nuclear (ITS, ETS) and plastid (matK, trnS-trnG and trnL-trnF) markers and ITS had the highest number of informative characters (Duarte et al. 2011, Carvalho-Sobrinho et al. 2016). In spite of having drawbacks related to potential paralogous copies (Álvarez 2003, Buckler et al. 1997), ITS can still play an important role in the investigation of species relationships if analysed carefully, for example identifying pseudogenes and assessing orthology in the case of intra-individual polymorphism (Bailey 2003, Feliner and Rosselló 2007).

Genomic DNA extraction was performed for 36 herbarium and silica-gel dried leaf samples using Qiagen DNeasy Plant Mini Kits following the manufacturer’s protocol, with the following changes: twice the volume of buffer AP1 in addition to a small amount of PVPP (polyvinyl polypyrrolidone) added at the lyse step followed by an incubation of 30 minutes; addition of 1uL of Riboshredder in the lysate solution followed by incubation at 37°C for 20 minutes; addition of twice the volume of buffer P3; and final elution in 46 uL of EB buffer run through the column twice to increase yield. Each 20 uL PCR amplification reaction contained 0.5 uL of template, 2 uL of dNTPs (2 mM), 2 uL of 10x reaction buffer, 1 uL of MgCl2 (50 mM), 0.65 uL of both forward primer and reverse primer solutions (10 uM), 0.1 uL of Taq polymerase, 4 uL of CES buffer and 9.1 uL of ddH2O. Amplification followed the same procedure described in Carvalho-Sobrinho et al. (2016). Samples were submitted to the Edinburgh Genomics laboratory at the University of Edinburgh for sequencing. For low quality sequences, we tested variations of the protocol (e.g. diminishing the amount of template in the PCR reaction or varying the sequencing primer). High quality sequences were recovered for 13 out of the 36 samples from which DNA was extracted for the ITS region.

All the inter-accession polymorphisms detected were validated visually by checking the electropherograms. Sequences were edited with Sequencher 5.4.1 (Gene Codes Corp., Ann Arbor, Michigan) and alignments were performed manually in Mesquite (Maddison and Maddison 2015). We investigated the potential presence of ITS pseudogenes by conducting phylogenetic analysis in separate partitions representing the 5.8S (conserved region) and the ITS 1 and ITS 2 regions (fast evolving regions). In pseudogenes, the conserved and the fast evolving regions are expected to show similar rate of evolution whereas in functional genes the conserved region is expected to have a lower rate of evolution compared to the fast evolving regions (Bailey et al. 2003).We assigned partitions by comparison with the annotated accession of *Pseudobombax tomentosum* (GenBank KM453206), and checked for differences in rates of evolution in the partition scheme proposed with PartitionFinder2 version 2.1.1 (Lanfear et al. 2017) using PhyML version 3.0 (Guindon et al. 2010) and the greedy search algorithm (Lanfear et al. 2012).

### Phylogenetic analysis and molecular dating

We implemented maximum likelihood (ML) and Bayesian Inference (BI) analysis. To determine best fitting model of sequence evolution for each of the three partitions, we used PartitionFinder2 version 2.1.1 (Lanfear et al. 2017) in the ML analysis and the reversible jump model selection (RB) implemented in BEAST2 version 2.5.1 (Bouckaert et al. 2019) under the Bayesian framework. IQ-TREE version 2.0.3 (Nguyen et al. 2015, Minh et al. 2020) was used to run the ML analysis with 1,000 bootstrap replicates and using the partition model option (-p) (Chernomor et al. 2016) with substitution models specified as follows: GTR+G for ITS1, K80+G for 5.8S and HKY+G for ITS2, as inferred with PartitionFinder2 (see above). BEAST2 version 2.5.1 (Bouckaert et al. 2019) was used to perform BI analysis and temporally calibrate the phylogeny. Different combinations of relaxed clock models (Uncorrelated Exponential Distribution - UCED and Uncorrelated Log Normal Distribution - UCLD (Drummond et al. 2006)) and tree priors (Yule and Birth-Death) were compared. Few studies have objectively contrasted the effect of different models in the divergence time estimation, and a poorly inferred time-calibrated phylogeny can have serious consequences for our understanding of diversification history of lineages (Louca and Pennell 2020). For example, different tree priors resulted in impressive differences in age estimation for cycads, with the Yule prior inferring ages three times older than the Birth-Death prior (Condamine et al. 2015). In Bayesian analysis, the most suitable model can be selected by comparing the Bayes Factor (BF). The BF is equal to the ratio of the Marginal Likelihood Estimate (MLE) of two models (BF=MLE1/MLE2) or to the difference of MLEs in log space (log(BF)=log(MLE1) – log(MLE2)). Positive values of BF would favour MLE1, and different values have different strengths. Values above five indicates that one model is significantly favoured over the other (Kass and Raftery 1995, Condamine et al. 2015), values above 20 indicate strong support and values above 150 overwhelming support. We estimated the Marginal Likelihood using the Nested Sampling algorithm (Skilling 2006) implemented in the NS package version 1.0.4 (Russel et al. 2019) for BEAST2 version 2.5.1 with 60 particles and 10,000 chain length. The NS package also calculates the standard deviation (SD) of the estimated Marginal Likelihood, which allows us to have confidence in the BF values calculated. For each combination of priors, we ran two independent runs of 10 million generations, sampled every 1,000 generations and visually inspected convergence of MCMC and ensured effective sample size > 200 for all parameters of each run using Tracer v1.7.1 (Rambaut et al. 2018). Resulting trees and log files from each run were combined using LogCombiner with a burn-in of 10% and the Maximum Clade Credibility Tree was summarized in TreeAnnotator with node heights as mean heights. We used r8s (Sanderson 2004) to implement the penalised likelihood method (Sanderson 2002) and calculate substitutions rates. We used the phylogram derived from the ML analysis as input and conducted a cross-validation analysis to find the best smoothing parameter.

We used the fossil flower of *Eriotheca prima* (Duarte 1974) from the middle to late Eocene (de Lima and Salard-Cheboldaeff 1981) of Brazil as a primary calibration for our BEAST2 analysis. The flower was identified as *Eriotheca* based on its small size (*Bombacopsis* and *Pachira* have larger flowers) and androecium organisation, which is a synapomorphy for the extant species of the genus (Robyns 1963, Duarte et al. 2011, Carvalho-Sobrinho et al. 2016). In previous studies, *Eriotheca* was resolved as sister to a clade comprising *Pseudobombax*, *Spirotheca, Ceiba*, and *Pochota fendleri* (Duarte et al. 2011, Carvalho-Sobrinho et al. 2016). Because the dating of this fossil is imprecise (middle to late Eocene: 33-56 million years old (Ma), we assigned the offset age of 33 Ma as a minimum age to the stem node (Renner 2005, Pennington et al. 2006) of *Eriotheca* (the crown node of the clade comprising *Eriotheca*, *Spirotheca*, *Pseudobombax*, *Pochotoa fendleri* and *Ceiba*), which is equivalent to the root node of the outgroups and ingroup of this study. We assigned a log-normal distribution with a mean of 1.542 and standard deviation of 1.5. This fossil calibration is conservative, with 95% of the prior distribution comprised between 33 and 47 Ma and thus the ages estimated here are considered minimum ages estimates. In order to explore the effects of using the medium and maximum ages of the *Eriotheca* fossil on phylogenetic age estimates, we also ran analyses assigning minimum ages of 47 Ma and 56 Ma to the *Eriotheca* stem (Figs. S1 and S2). We followed the dates on the Geologic Time Scale v. 5.0 (Gradstein et al. 2012).

### Phylogenetic signal test

We tested for strength of phylogenetic signal for the binary traits related to ecology (rain vs dry forests) and geography (Central and North America vs South America) using the *D value* proposed by Fritz and Purvis (2010), and implemented using the Caper package (v. 1.0.1) (Orme 2013) in R, with 5,000 permutations. Under a null model of Brownian motion evolution of a binary trait, *D* has an expected value of 0. A negative *D* value indicates a strongly clustered phylogenetic pattern for a given binary trait (perhaps due to some process of evolutionary constraint), a value of one indicates a completely random pattern with respect to the phylogeny (i.e. no correlation between phylogeny and the trait at all) and values above one indicate an overdispersed phylogenetic pattern (perhaps due to divergent selection). We assigned species to ecological and geographical categories following Gibbs and Semir (2003) (Table S1). Despite occurring mainly in rain forests, *Ceiba speciosa* is also recorded in dry semi-deciduous woodland (Figure 2). To check for possible bias in the ecological affinity of *C. speciosa*, we also conducted a phylogenetic signal analysis assigning this species to dry forest, observing no difference when comparing to the analysis run assigning the species to rain forest.

## RESULTS

The total length of the aligned sequences was 814 nucleotides, of which 283 were variable and 178 (22%) were parsimony-informative characters. The ML and BI trees showed congruent topologies (Figs. 3 and 4). *Ceiba* was strongly supported as monophyletic, with posterior probability (pp) = 1 and bootstrap value = 100 and was recovered as sister to *Pseudobombax* (Fig. 3). The UCLD and UCED clock models and Yule and Birth-Death tree models inferred similar crown and stem ages with overlapping credibility intervals (95% Highest Posterior Density (HDP), Table 1). For the UCLD clock prior, the Bayes Factor value of 6.15 support the Yule tree prior as the most suitable tree model. For the UCED clock prior, the Bayes Factor of 1.15 indicates that the neither tree model is favoured (Table 1). Therefore, results shown onwards for Bayesian analysis are those inferred using the UCLD clock model and the Yule tree model. Using the 33 Ma fossil calibration, the stem node age of *Ceiba* is 21.1 (14.7-27.1 [95% HPD]) Ma, the crown node age is 12.7 (8.2-17.6 [95% HPD]) Ma (Table 1, Fig. 4), and substitution rates estimated in 1.592 ×10^−9^ substitutions per site per year (s/s/y). *Ceiba* shows slow substitution rates for ITS when compared to other tropical tree species. For example, for the rain forest tree genus *Inga* (Leguminosae), substitution rates have been estimated in 2.34 × 10^−9^ s/s/y (Richardson et al. 2001) and in 7.1-7.9 × 10^−9^ s/s/y (Lavin 2006), and for the dry forest genus *Coursetia* (Leguminosae) in 5.0-8.2 × 10^−9^ (Lavin 2006).

**Table 1.** Absolute ages estimate for ten nodes under different tree (Yule and Birth-Death) and clock priors (UCLD – Uncorrelated Lognormal Distribution and UCED – Uncorrelated Exponential Distribution). Ages are reported in million years as mean ages followed by the 95% Highest Posterior Density (HDP) as a result of the combined independent runs for each tree and clock priors. MLE (SD): marginal likelihood estimated followed by standard deviation; BF: Bayes factor calculated as the difference between the MLE of the Yule and the Birth-Death prior for each clock prior UCLD and UCED. Values above five indicates that one model is significantly favoured over the other.

| Clock prior | UCLD |  | UCED |  |
| --- | --- | --- | --- | --- |
|  | Yule | Birth-death | Yule | Birth-death |
| **MLE (SD)** | -3844.31 (1.85) | -3838.15 (1.79) | -3838.74 (1.88) | -3837.62 (1.76) |
| **BF** | 6.15 |  | 1.15 |  |
| ***Ceiba* crown** | 12.7 (8.2-17.6) | 10.9 (6.5-15.7) | 13.5 (8.6-18.8) | 11.5 (6.1-17.4) |
| ***Ceiba* stem** | 21.1 (14.7-27.1) | 20.3 (13.5-26.9) | 21.0 (14.4-27.5) | 19.1 (11.3-26.9) |
| **SDTF SA crown** | 8.6 (5.0-12.5) | 7.0 (3.6-10.7) | 9.6 (5.6-14.1) | 7.8 (3.8-12.5) |
| **SDTF SA stem** | 11.2 (7.2-15.6) | 9.5 (5.6-13.9) | 12.0 (7.5-16.7) | 10.0 (5.2-15.4) |
| ***C. aesculifolia* crown** | 1.5 (0.2-3.2) | 1.2 (0.2-2.3) | 1.7 (0.3-3.9) | 1.3 (0.2-2.9) |
| ***C. aesculifolia* stem** | 6.3 (3.1-9.8) | 5.1 (2.1-8.3) | 7.6 (3.7-12.0) | 4.7 (1.7-8.3) |
| ***C. samauma* crown** | 0.4 (0.0001-1.4) | 0.3 (0.0001-1.0) | 0.5 (0.0001-1.7) | 0.3 (0.0001-1.2) |
| ***C. samauma* stem** | 6.3 (3.1-9.8) | 5.1 (2.1-8.3) | 6.3 (2.6-10.2) | 6.1 (2.4-10.3) |
| ***C. pentandra* crown** | 3.9 (1.6-6.5) | 3.0 (1.1-5.2) | 5.5 (2.3-9.2) | 4.2 (1.5-7.5) |
| ***C. pentandra* stem** | 12.7 (8.2-17.6) | 10.9 (6.5-15.7) | 13.5 (8.6-18.9) | 11.5 (6.1-17.4) |

**Figure 3.**
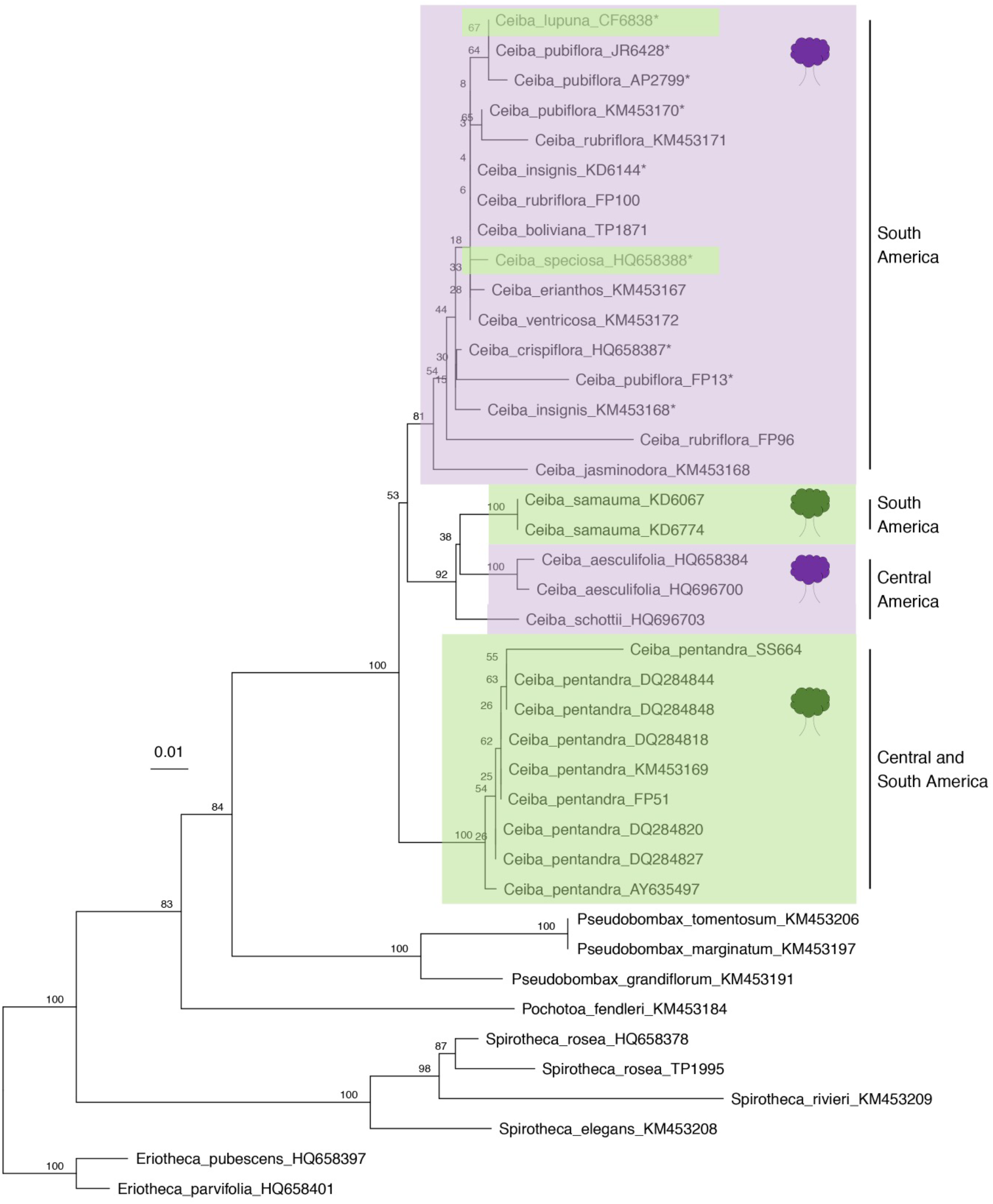
Maximum likelihood phylogram derived from analysis of nuclear ribosomal ITS sequence data sets for 14 species of *Ceiba*. Species with asterisk belong to the *Ceiba insignis* species aggregate. Values above branches represent bootstrap values for internal nodes. Tree symbols in front of accessions represent species occurring in SDTF (purple) and rain forests (green).

**Figure 4.**
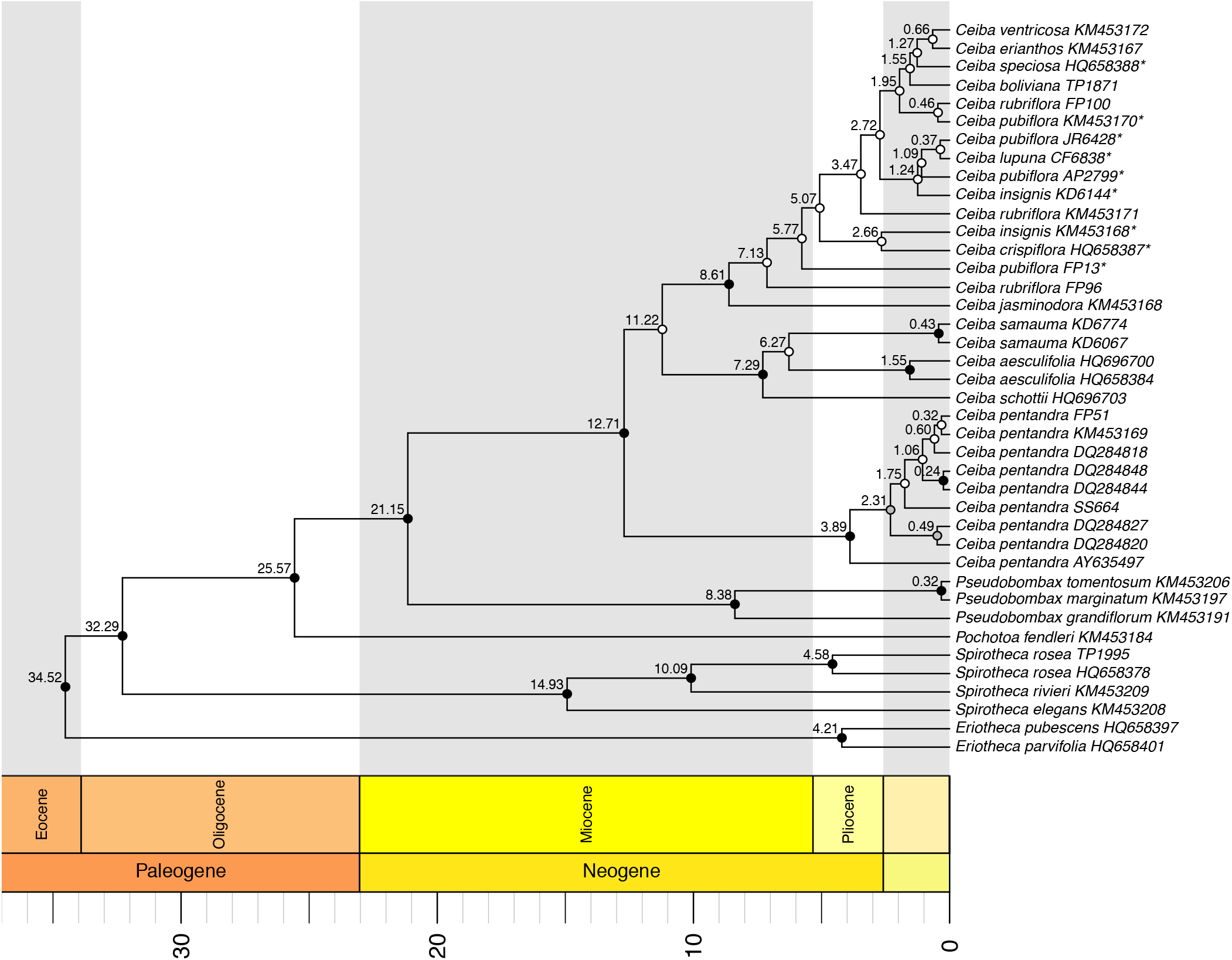
Maximum clade credibility tree resulting from BEAST2 analysis of nuclear ribosomal ITS sequence data sets for 14 species of *Ceiba*, 33 Ma offset calibration, using Yule tree prior and the Uncorrelated Lognormal Distribution clock model. Values above branches represent nodes ages reported in million years as mean ages. Circles represent posterior probabilities for internal nodes: black >= 0.95; grey < 0.95 and >= 0.75, and white < 0.75.

*Ceiba* comprises three main clades: (i) a rain forest lineage of the three accessions of *C. pentandra*, which are strongly supported as monophyletic [posterior probability (pp) = 1 and bootstrap value = 100] and sister to the remaining species and with stem node age of 12,7 (8,2-17,6 [95% HPD]) Ma and crown node age of 3,9 (1.6–6,5 [95% HPD]) Ma.; (ii) a highly supported clade [posterior probability (pp) = 1 and bootstrap value = 92] composed of *C. schottii* and *C. aesculifolia* from Central American and Mexican SDTF plus two accessions of *C. samauma* from inter-Andean valleys in Peru, with stem node age of 11,2 (7,2-15,6 [95% HPD]) Ma and crown age of 7,3 (3,7-11.0 [95% HPD]) Ma; and (iii) a highly supported [posterior probability (pp) = 0,99 and bootstrap value = 81] South American SDTF clade including 10 species showing little sequence variation, with stem node age of 11,2 (7,2–15,6 [95% HPD]) Ma and crown node of 8,6 (5,0–12,5 [95% HPD]) Ma. Within this South American clade, *C. rubriflora*, *C. pubiflora* and *C. insignis*, which were represented by multiple accessions, were resolved as monophyletic. The South American clade represents SDTF species, except for *C. lupuna*, a species with a distribution restricted to rain forest (Figs. 3 and 4). *Ceiba aesculifolia* was the only SDTF species recovered as monophyletic with stem node age of 6.3 (3.1–9.8 [95% HPD]) Ma and crown node of 1.5 (0.2–3.2 [95% HPD]) Ma.

The *D* test shows significant phylogenetic signal for both ecological preference (*D* = 0.1247542, P (D=1) = 0.001, P (D=0) = 0.3422) and geographical occurrence (*D*= 0.2204991, P (D=1) = 0.0176, P (D=0) = 0.3152). Both *D* values are statistically indistinguishable from zero, which indicates that closely related species are more likely to show the same ecological preference or geographical occurrence, as expected under a Brownian model of evolution, whereby there would be a constant rate of state switching over time and any given lineage is more likely to stay within the same biome and geographic region per unit time than to switch to the alternative biome or geographic region.

The phylogeny supports the monophyly of the two sections of the genus, *Ceiba* and *Campylanthera*, which are based on pollen and staminal appendages characters. However, it does not support monophyly of the ‘insignis’ species complex (Gibbs and Semir 2003). This species aggregate includes seven species (*C. pubiflora*, *C. chodatii*, *C. insignis*, *C. ventricosa*, *C. lupuna*, *C. speciosa* and *C. crispiflora*, indicated with an asterisk in Figs. 3 and 4) characterized by their entire staminal tube terminating in a collar of anthers, with the exception of *C. pubiflora* which has free stamens.

## DISCUSSION

### Geographic and ecological structure

The D test shows clear phylogenetic signal for ecological preference and geographic phylogenetic structure (i.e., clear Central and South American clades) in *Ceiba.* However, our data suggest multiple shifts from dry to rain forests within *Ceiba* (Figs. 3 and 4) because rain forest species are nested within the two dry forest clades. For example, the two accessions representing *C. samauma*, occurring in rain and riverine forest in South America, are nested within the Central American and Mexican clade and the rain forest species *C. lupuna* and *C. speciosa*, are nested within the South American SDTF clade.

### Biome-specific differences in the nature of species and their diversification trajectories

Our results show young crown and stem ages for species in the SDTF clade. Rain forest species such as *Ceiba pentandra* and *C. samauma* show patterns of long stems with shallow crown groups (Table 1). These patterns contrast to previous studies of individual SDTF species that showed them to be older, with stem ages of 5-10my (e.g., Pennington et al. 2010, de Queiroz & Lavin 2011), and runs contrary to the prediction of Pennington and Lavin (2016) that rain forest species might, on average, tend to have more recent origins.

The stem age of *C. pentandra* is estimated at 12.7 Ma (Table 1). The long stem and shallow crown suggest this is an old rain forest lineage with low levels of extant genetic diversity populations. Likewise, *C. samauma* was recovered as monophyletic, and has a crown node age estimated as 0.4 Ma and a stem node age of 6,3 Ma. Both species therefore contrast with the suggested predominant patterns for rain forest species. Our result, recovering *C. pentandra* as monophyletic, with low sequence divergence amongst accessions, is consistent with that of Dick et al. (2007) who showed *C. pentandra* to have extremely weak phylogeographical structure based on ITS and chloroplast *psb*B- *psb*F for 51 individuals. In addition to that, the disjunct distribution of this species in Africa was demonstrated to be due to relatively recent long distance dispersal because of low genetic divergence of the African populations.

Within the two predominantly SDTF clades, there is little evidence for old lineages representing morphologically recognized species with long stems and monophyletic crown groups, as predicted by Pennington and Lavin (2016). The crown age of the South American SDTF clade, containing 10 species, is estimated at 8.6 Ma, and the Mexican SDTF clade, containing 2 species, is estimated at 7.9 Ma with a stem age for both estimated at 11.2 Ma. Only one species from SDTF were recovered as monophyletic, *Ceiba aesculifolia* with a crown age estimated at 1.5 Ma and stem age at 6.3 Ma. Even when assigning a minimum age of 56 Ma to the *Eriotheca* stem, the same pattern is observed (Fig. S2). The crown age of the South American SDTF clade is estimated at 14.9 Ma and the Mexican SDTF clade, containing two species, is estimated at 12.6 Ma with a stem age for both estimated at 19.5 Ma. The crown age of *Ceiba aesculifolia* was estimated at 2.7 Ma and stem age at 10.79 Ma (Fig. S2).

The lack of resolution among the dry forest accessions, with most species being recovered as non-monophyletic, indicates absence of intraspecific coalescence for the ITS locus.

Explanations for this include incomplete lineage sorting after speciation events, paralogous gene copies, inaccurate species delimitation and/or hybridization followed by introgression (Naciri and Linder 2015, Pennington and Lavin 2016). We eliminated sequences with possible paralogues by visual inspection of the electropherograms and by comparing substitution rates along branch lengths following Bailey et al. (2003). Some species of *Ceiba* are hypothesised to be interfertile and hybridise (Gibbs and Semir 2003), especially within the *insignis* species aggregate. However, it is also suggested that those species diverge in time of anthesis and pollinator type as well, and we have seen no evidence of putative hybrids in the field (Pezzini, pers. obs.). Eight out of the ten species within the South American SDTF clade are from Brazil and of these, four are distributed in the *Caatinga*, the largest area of SDTF in the Neotropics (700,000km2) (Silva de Miranda et al. 2018). *Ceiba* species such as *C. pubiflora* are often widespread (Fig. 2) and abundant (Lima et al. 2010). Taken together, this evidence suggests that the non-monophyly of *Ceiba* species found in SDTF such as *C. pubiflora* may be a reflection of large effective population sizes and hence a longer time to coalescence (Naciri and Linder 2015, Pennington and Lavin 2016), rather than due to hybridisation or ITS paralogy.

Our study illustrates that the general patterns of species age, monophyly and geographical structure reported for species belonging to the Leguminosae family and endemic to SDTF (Pennington and Lavin 2016) are not shared by one of the most characteristic SDTF tree genera and suggests that further phylogenetic studies of unrelated groups are required.

### Taxonomic implications

Our data support: (i) the circumscription of *Chorisia* within *Ceiba*, as proposed by Gibbs et al. (1988), Ravenna (1998) and Gibbs and Semir (2003) and confirmed by recent molecular studies (Carvalho-Sobrinho et al. 2016); (ii) the non-monophyly of the *C. insignis* aggregate species proposed by Gibbs & Semir (2003). Our data suggest that *C. boliviana*, *C. erianthos* and *C. rubriflora*, not included by Gibbs and Semir (2003) are also part of the *insignis* clade (Fig. 3). It is suggested that those species are interfertile but also diverge in time of anthesis and pollinator type (Gibbs and Semir 2003). Five of the seven species within this complex are restricted to the SDTF patches of South America, while *C. speciosa* is widespread and *C. lupuna* occurs in riverine rain forests in the Peruvian and Brazilian Amazon (Fig. 2); and (iii) the monophyly of the section *Campylanthera* (Gibbs and Semir 2003) that includes the Central American species *C. aesculifolia*, *C. schottii* and the widespread *C. samauma*.

## ACKNOWLEDGMENTS

This paper is part of the PhD thesis of F.P. F.P. acknowledges the support of the Brazilian National Research Council (CNPq) for funding her PhD at the University of Edinburgh and the Royal Botanic Garden Edinburgh under the Science without Borders programme (process 206954/2014-0 - GDE). Fieldwork was partially supported by Fundação de Amparo à Pesquisa do Estado da Bahia (FAPESB - process APP0006/2011). F.P. thanks F.C. Diniz for fieldwork collaboration, A. Griffths for help with the phylogenetic signal analysis and E. Koenen for insightful discussions. The authors are thankful to the reviewers who helped improve the manuscript.

## Author contributions

F.F.P. and R.T.P conceived the original idea; F.F.P. executed fieldwork, data collection, laboratory work and data analysis with input from K.G.D; F.F.P. led the writing of the manuscript with input, comments and review from K.G.D., J.G.C.-S., C.A.K., J.A.N. L.P.Q. and R.T.P.

## DATA AVAILABILITY STATEMENT

### Data Availability

The data used in this study are archived at Genbank (accession numbers XXX – numbers will be inserted on final accepted version). Scripts for analysis conducted here and for making figures are available in F.F.P.’s GitHub page (https://github.com/fpezzini).

## Supplementary Materials

The following materials are available as part of the online article: Supplementary Table 1. Collection details of each accession and ecological preference for the species of *Ceiba*; Figure S1. Maximum clade credibility tree resulting from BEAST2 analysis of nuclear ribosomal ITS sequence data sets for 14 species of *Ceiba* for 47 Ma offset calibration, using Yule tree prior and the Uncorrelated Lognormal Distribution clock model; Figure S2. Maximum clade credibility tree resulting from BEAST2 analysis of nuclear ribosomal ITS sequence data sets for 14 species of *Ceiba* for 56 Ma offset calibration, using Yule tree prior and the Uncorrelated Lognormal Distribution clock model.

## Supplementary Material

**Supplementary Table 1.** Collection details of each accession and ecological preference for the species of *Ceiba.*

| n | Accession | Species | Locality | Country | GenBank accession | Collector name + number (Herbarium) | Ecological preference |
| --- | --- | --- | --- | --- | --- | --- | --- |
| 1 | HQ658384 | *Ceiba aesculifolia* | - | - | HQ658384 | Fairchild Botanical Gardens acc. no. 83301 | SDTF |
| 2 | HQ696700 | *Ceiba aesculifolia* accession2 | - | - | HQ696700 | Fairchild Botanical Gardens acc. no. X-2-206 | SDTF |
| 3 | TP1871 | *Ceiba boliviana* | Huancavelica | Peru | in process of submission | Pennington, R. Toby (E) | SDTF |
| 4 | HQ658387 | *Ceiba crispiflora* | - | - | HQ658387 | Pacific Tropical Garden acc. no. 750726001 | rain forests |
| 5 | KM453167 | *Ceiba erianthos* | Bahia | Brazil | KM453167 | E.R.Souza 710 (HUEFS) | SDTF |
| 6 | KD6144 | *Ceiba insignis* | Pucacaca | Peru | in process of submission | Dexter, K.G. (E) | SDTF |
| 7 | KM488629 | *Ceiba insignis* | Cajamarca | Peru | KM488629 | J. Campos & P. López 4953 (US) | SDTF |
| 8 | KM453168 | *Ceiba jasminodora* | Rio de Janeiro | Brazil | KM453168 | Carvalho-Sobrinho 3070 (HUEFS) | SDTF |
| 9 | CF6838 | *Ceiba lupuna* | Acre | Brazil | in process of submission | Daly, D.C. 6838 (E) | rain forests |
| 10 | FP51 | *Ceiba pentandra* | Piauí | Brazil | in process of submission | Pezzini, F.F. 51 (HUEFS) | rain forests |
| 11 | SS664 | *Ceiba pentandra* | Minas | Brazil | in process of submission | Sant'ana, S.C. (E) | rain forests |
| 12 | KM453169 | *Ceiba pentandra* | - | - | KM453169 | Carvalho-Sobrinho s.n. (HUEFS) | rain forests |
| 13 | DQ284818 | *Ceiba pentandra* | Camuy | Puerto Rico | DQ284818 | - | rain forests |
| 14 | DQ284848 | *Ceiba pentandra* | Kourou | French Guiana | DQ284848 | - | rain forests |
| 15 | DQ284844 | *Ceiba pentandra* | Castanhal Veadó (Rio Trombetas) | Brazil | DQ284844 | - | rain forests |
| 16 | DQ284827 | *Ceiba pentandra* | Borbon | Ecuador | DQ284827 | - | rain forests |
| 17 | DQ284820 | *Ceiba pentandra* | Morelos | Mexico | DQ284820 | - | rain forests |
| 18 | AY635497 | *Ceiba pentandra* | Jobero | Panama | AY635497 | - | rain forests |
| 19 | FP13 | *Ceiba pubiflora* | Bahia | Brazil | in process of submission | Pezzini, F.F. 13 (HUEFS) | SDTF |
| 20 | JR6428 | *Ceiba pubiflora* | Minas | Brazil | in process of submission | Ratter, J. (E) | SDTF |
| 21 | AP2799 | *Ceiba pubiflora* | Minas | Brazil | in process of submission | A. Pott (E) | SDTF |
| 22 | KM453170 | *Ceiba pubiflora* | Bahia | Brazil | KM453170 | Carvalho-Sobrinho 3066 (HUEFS) | SDTF |
| 23 | FP96 | *Ceiba rubriflora* | Minas | Brazil | in process of submission | Pezzini, F.F. 96 (HUEFS) | SDTF |
| 24 | FP100 | *Ceiba rubriflora* | Minas | Brazil | in process of submission | Pezzini, F.F. 100(HUEFS) | SDTF |
| 25 | KM453171 | *Ceiba rubriflora* | Bahia | Brazil | KM453171 | Carvalho-Sobrinho 574 (HUEFS) | SDTF |
| 26 | KD6067 | *Ceiba samauma* | Buenos Aires | Peru | in process of submission | Dexter, K.G. (E) | rain forests |
| 27 | KD6774 | *Ceiba samauma* | Echarate | Peru | in process of submission | Dexter, K.G. (E) | rain forests |
| 28 | HQ696703 | *Ceiba schottii* | - | - | HQ696703 | Fairchild Botanical Gardens acc. no. 83302 | SDTF |
| 29 | HQ658388 | *Ceiba speciosa* | - | Brazil | HQ658388 | W.S. Alverson s.n. (WIS) | rain forests |
| 30 | KM453172 | *Ceiba ventricosa* | Bahia | Brazil | KM453172 | Carvalho-Sobrinho 3124 (HUEFS) | SDTF |
| 31 | HQ658401 | *Eriotheca parvifolia* | - | - | HQ658401 | M.C. Duarte 109 (SP) | - |
| 32 | HQ658397 | *Eriotheca pubescens* | - | - | HQ658397 | M.C. Duarte 115 (SP) | - |
| 33 | KM453184 | *Pochota fendleri* | - | - | KM453184 | P.E. Kaminski s.n (HUEFS) | - |
| 34 | KM453191 | *Pseudobombax grandiflorum* | - | - | KM453191 | Carvalho-Sobrinho 2946 HUEFS | - |
| 35 | KM453197 | *Pseudobombax marginatum* | - | - | KM453197 | L.P. Queiroz 14753 (HUEFS) | - |
| 36 | KM453206 | *Pseudobombax tomentosum* | - | - | KM453206 | Carvalho-Sobrinho 2874 HUEFS | - |
| 37 | KM453208 | *Spirotheca elegans* | - | - | KM453208 | Carvalho-Sobrinho 2964 HUEFS | - |
| 38 | KM453209 | *Spirotheca rivieri* | - | - | KM453209 | Carvalho-Sobrinho s.n. HUEFS | - |
| 39 | TP1995 | *Spirotheca rosea* | Junin | Peru | in process of submission | Pennington, R. Toby (E) | - |
| 40 | HQ658378 | *Spirotheca rosea* | - | - | HQ658378 | W.S. Alverson 2185 (WIS) | - |

**Supplementary Figure 1.**
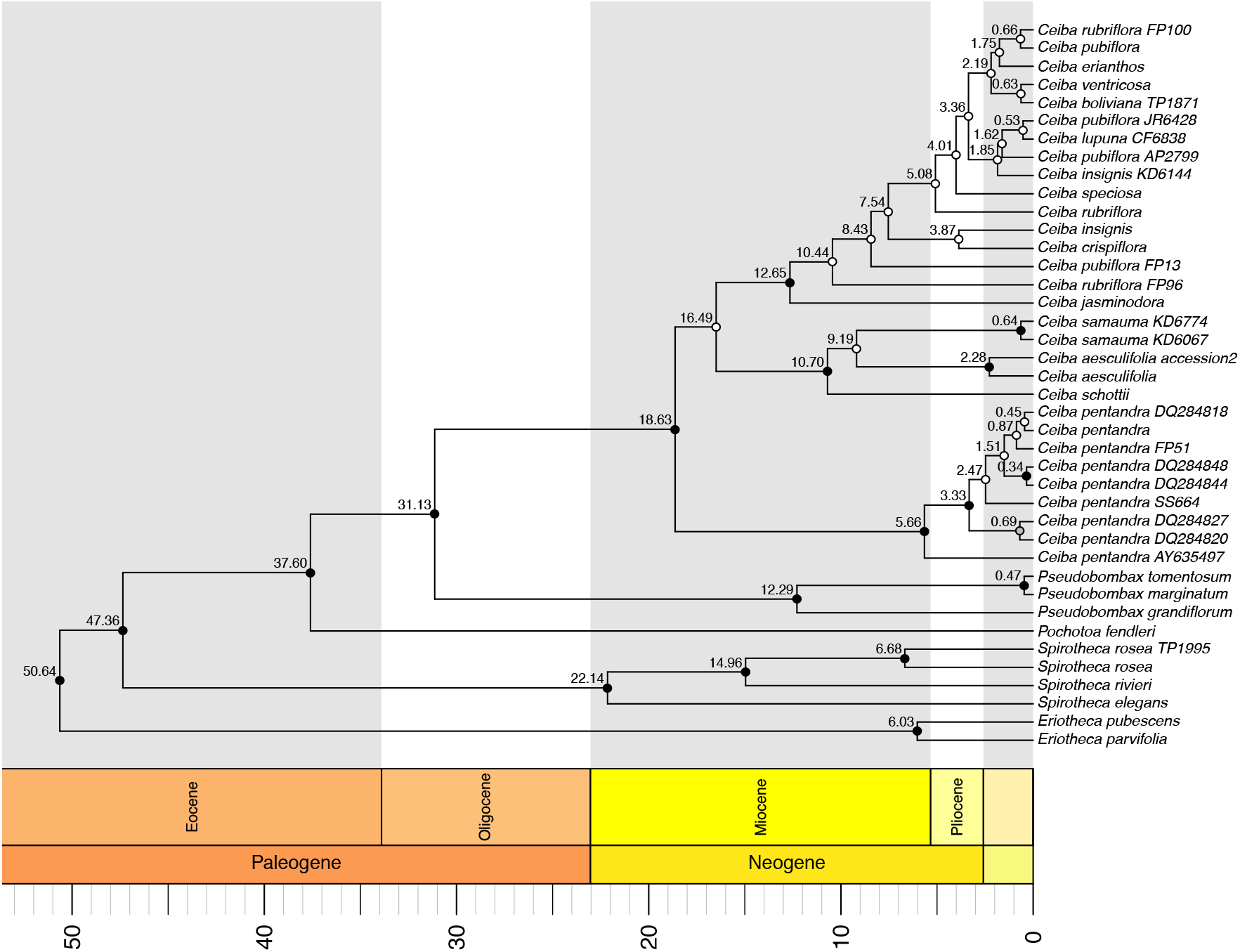
Maximum clade credibility tree resulting from BEAST2 analysis of nuclear ribosomal ITS sequence data sets for 14 species of *Ceiba* and 47 Ma offset calibration, using Yule tree prior and the Uncorrelated Lognormal Distribution clock model. Values above branches represent nodes ages reported in million years as mean ages. Circles represent posterior probabilities for internal nodes: black >= 0.95; grey < 0.95 and >= 0.75, and white < 0.75.

**Supplementary Figure 2.**
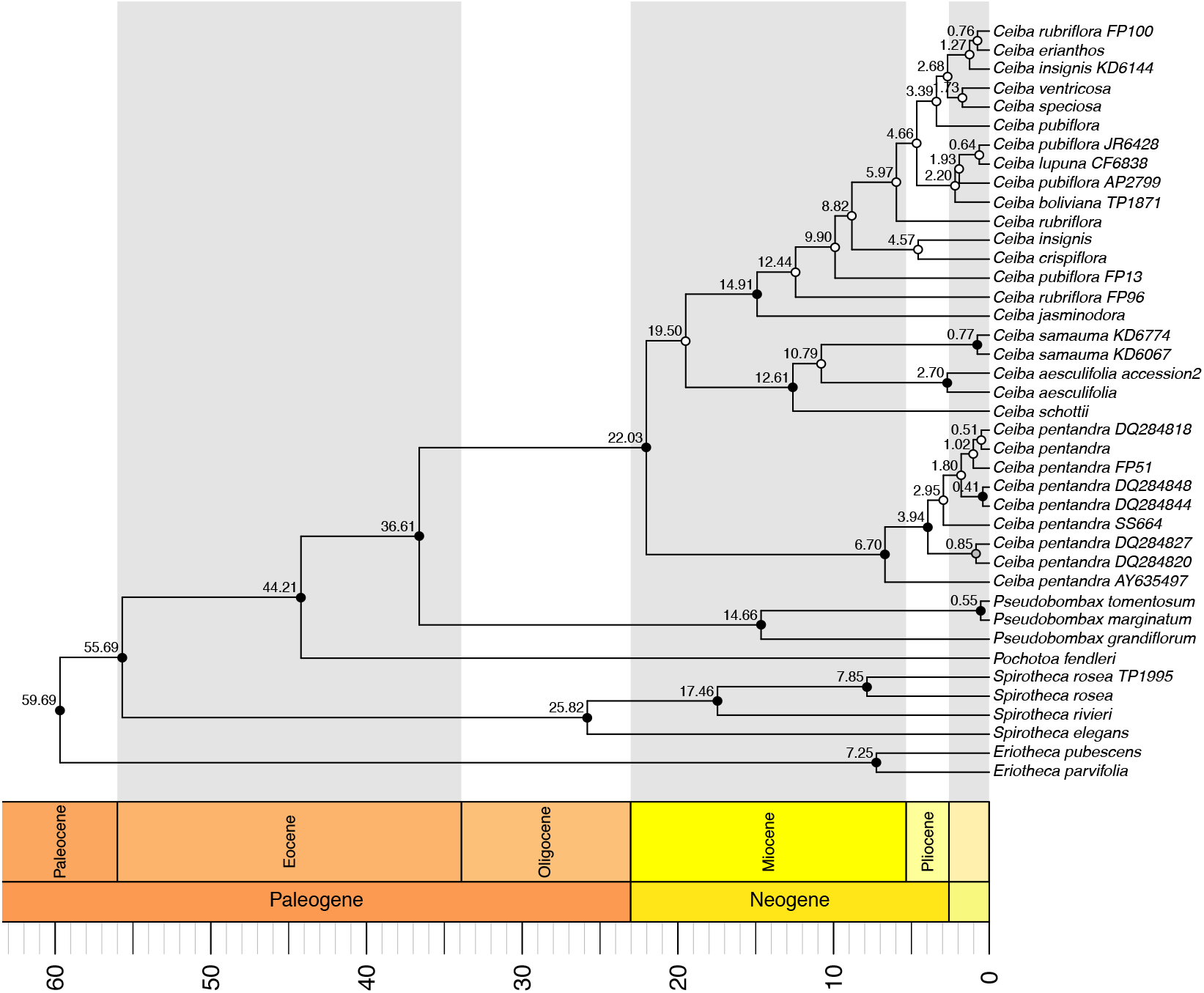
Maximum clade credibility tree resulting from BEAST2 analysis of nuclear ribosomal ITS sequence data sets for 14 species of *Ceiba* and 56 Ma offset calibration, using Yule tree prior and the Uncorrelated Lognormal Distribution clock model. Values above branches represent nodes ages reported in million years as mean ages. Circles represent posterior probabilities for internal nodes: black >= 0.95; grey < 0.95 and >= 0.75, and white < 0.75.

